# TP promotes malignant progression in hepatocellular carcinoma through pentose Warburg effect

**DOI:** 10.1101/386706

**Authors:** Qiang Zhang, Yuan Qin, Jianmin Zhao, Yuanhao Tang, Xuejiao Hu, Weilong Zhong, Mimi Li, Shumin Zong, Meng Li, Honglian Tao, Zhen Zhang, Shuang Chen, Huijuan Liu, Lan Yang, Honggang Zhou, Yanrong Liu, Tao Sun, Cheng Yang

**Affiliations:** State Key Laboratory of Medicine Chemical Biology and College of Pharmacy, Nankai University, Tianjin, China; Tianjin Key Laboratory of Molecular Drug Research, Tianjin International Joint Academy of Biomedicine, Tianjin, China; Department of Pathology, Hospital of Shun Yi District, Beijing, China; College of Life Science, Nankai University, Tianjin, China

**Keywords:** Twist1, thymidine phosphorylase (TP), pentose Warburg effect, vasculogenic mimicry (VM)

## Abstract

Tumor progression is dependent on metabolic reprogramming. Metastasis and vasculogenic mimicry (VM) are typical tumor progression. The relationship of metastasis, VM and metabolic reprogramming is not clear. In this study, we identified the novel role of Twist1, a VM regulator, in the transcriptional regulation of the expression of thymidine phosphorylase (TP). We demonstrated that TP promoted extracellular thymidine metabolization into ATP and amino acids through pentose Warburg effect by coupling the pentose phosphate pathway and glycolysis. Moreover, Twist1 relied on TP-induced metabolic reprogramming to promote hepatocellular carcinoma (HCC) metastasis and VM formation mediated by VE-Cad, VEGFR1, and VEGFR2 in vitro and in vivo. TP inhibitor tipiracil reduced promotion effect of TP enzyme activity on HCC VM formation and metastasis. Our findings demonstrate that TP, transcriptionally activated by Twist1, promotes HCC VM formation and metastasis through pentose Warburg effect, contributing to tumor progression.

## Introduction

Hepatocellular carcinoma (HCC) is a leading cause of death from cancer worldwide, and HCC deaths largely result from vasculogenic mimicry (VM) formation and metastasis(Cao et al, 2013; Siegel et al, 2018; Uchino et al, 2011). Such malignant progression of HCC is a response to the deterioration of the local tumor microenvironment. Blood supply is required to sustain growth and metastasis. VM is a de novo microvascular channel formation by aggressive cancer cells that enables fluid transport from leaky vessels(Maniotis et al, 1999). The pathways involved in VM share components with stemness and epithelia–mesenchymal transition (EMT), key attributes that promote tumor metastasis(Kirschmann et al, 2012; Qiao et al, 2015). However, the mechanism by which tumor cells trigger VM formation is unclear.

Under a deteriorated local tumor microenvironment, tumor cells are forced to reprogram cellular metabolism(DeBerardinis et al, 2008). An obvious change in metabolism is the Warburg effect, where tumor cells mainly use glycolysis to generate energy even under aerobic conditions(Vander Heiden et al, 2009). This metabolic reprogramming eliminates the threat of hypoxia to the survival of tumor cells. Under a nutrient-poor environment, tumor cells may preferentially utilize glutamine as a source of nutrients(Pavlova et al, 2018). Moreover, tumor cells can use other carbon sources, such as lactate, serine, and glycine, as fuels(Faubert et al, 2017; Hui et al, 2017; Maddocks et al, 2017). By inducing paracancerous tissue cell autophagy, starved cancer cells can obtain fuel from extracellular sources(Sousa et al, 2016). Remarkably, distant metastases are dependent on the pentose phosphate pathway for reprogramming malignant gene expression and phenotype(McDonald et al, 2017). These metabolic reprogrammings could prevent tumor cells from surviving stress before VM formation. However, whether other metabolic reprogramming is involved in tumor malignant progression before VM formation and whether this metabolic reprogramming is related to VM formation and metastasis is unclear. Thus, further explorations are required.

Twist1 is a key transcription factor that induces EMT and VM by upregulating VE-Cadherin expression(Sun et al, 2010). In this study, we found that Twist1 also transcriptionally promotes the expression of thymidine phosphorylase (TP), also known as platelet-derived endothelial cell growth factor(Furukawa et al, 1992). When tumor vascular supply is occluded, TP exhibits a high expression under a hypoxic and low-pH environment(Griffiths et al, 1997). As a phosphorylase, TP catalyzes thymidine to deoxyribose-1-phosphate (dR-1-P), which can then be converted to dR-5-P, glyceraldehyde-3-phosphate (G-3-P), or deoxyribose(Bijnsdorp et al, 2010). TP promotes endothelium-dependent angiogenesis in endothelial cells(Miyadera et al, 1995). In a previous study, we demonstrated that TP promotes metastasis and serves as a poor prognostic marker in HCC(Zhang et al, 2017). In the present study, we explored whether TP upregulation affects the metabolic reprogramming of HCC and whether the transcriptional pattern of Twist1–TP could further contribute to VM formation in HCC.

In this study, we demonstrated that TP stimulates pentose Warburg effect by coupling the pentose phosphate pathway and glycolysis. The ATP and amino acids generated by the pentose Warburg effect further promote HCC VM formation and metastasis. Consequently, we revealed that TP, transcriptionally activated by Twist1, promotes HCC VM formation through pentose Warburg effect, contributing to tumor progression.

## Results

### Twist1 promotes TP transcription via binding to the conserved motifs in the promoter region

The candidate co-expression genes with Twist1 from the TCGA database were screened, and eventually 13 genes were significantly correlated with Twist1 (Figure 1A). ChIP results were used to screen the transcriptional targets of Twist1. We defined genes by binding peaks in the promoter region. The data revealed that 259 genes were targeted by Twist1 (Figure 1B, green circle). By Venn analysis, we identified TP in both predicted gene sets (Figure 1B). Further analysis of the LIHC data downloaded from the TCGA database showed that TP gene was more highly expressed in primary HCC samples compared with the normal liver samples (Figure 1C). To illustrate the malignant progression effects of TP in HCC, we grouped the clinical tissue samples based on clinical parameters. The Kaplan–Meier survival analysis results showed that high TP expression significantly affected the survival prognosis of HCC patients with clinical parameters of stage ≥ II, metastasis, or CEA content > 3.4 ng/mL, AFP content > 400 ng/mL, or ALT content > 38U/L (Figure S1).

**Figure 1.**
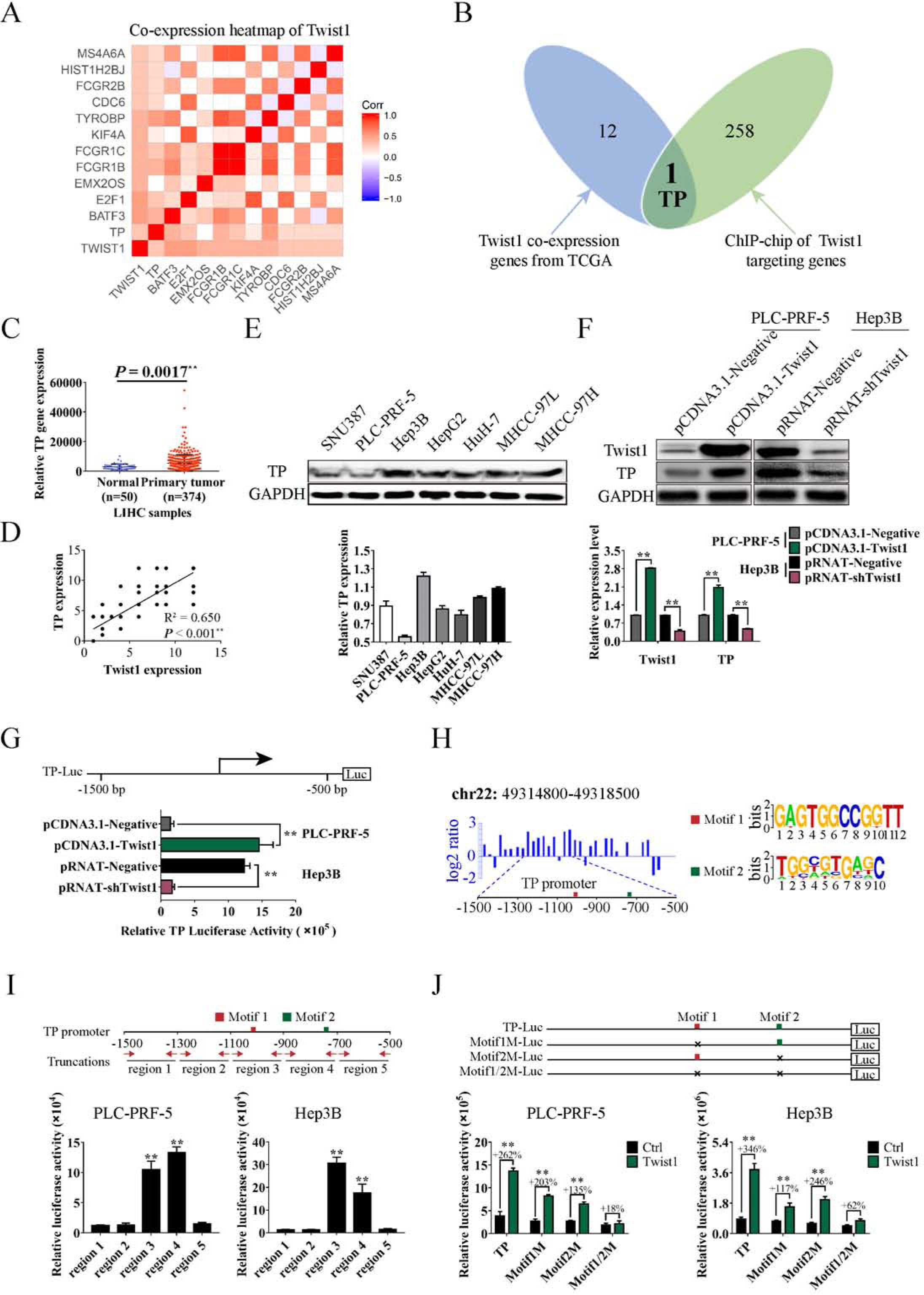
Twist1 promotes TP gene transcription via binding to the conserved motifs in the promoter region. **A.** Heat map of Twist1 co-expression genes downloaded from the TCGA database. **B.** Venn analysis to identify Twist1 target genes in human HCC. The blue diagram represents Twist1 co-expression genes from TCGA, and the green diagram represents Twist1 targeting genes by ChIP-seq. **C.** Expression analysis of TP in LIHC samples. **D.** Correlation analysis of Twist1 and TP expression in clinical HCC specimens. **E.** Background expression analysis of TP in seven HCC cell lines. **F.** Western blot was used to analyze the TP protein levels influenced by Twist1 in PLC-PRF-5 and Hep3B cell lines. **G.** Luciferase assay was performed to detect the TP gene transcription mediated by Twist1. **H.** Diagram of the ChIP-seq binding peaks related to the TP promoter. Binding site analysis showed two individual conserved motifs within the promoter of TP. **I.** Luciferase assay results of Twist1 to truncated TP promoter regions. **J.** Luciferase assay results of Twist1 to mutated TP promoter regions. (mean ± SD; n = 3 in triplicate; ^**^P < 0.01)

To further prove the transcriptional activation of TP by Twist1, we analyzed the correlation between Twist1 expression and TP expression in HCC samples. Results showed a significant correlation between these two proteins (Figure 1D). Seven HCC cells were chosen to detect the background expression levels of TP. PLC-PRF-5 and Hep3B were separately used as TP low-expression and high-expression cell line models for subsequent experiments (Figure 1E). Western blot results showed that upregulated Twist1 expression in the PLC-PRF-5 cells significantly promoted TP expression, and TP expression was significantly reduced when Twist1 was knocked down in the Hep3B cells (Figure 1F). Furthermore, the luciferase assay proved that ectopic overexpression of Twist1 enhanced the transcriptional activation of TP in the PLC-PRF-5 cells, whereas Twist1 downregulation reduced the activity in the Hep3B cells (Figure 1G). According to the peak regions from the ChIP-seq assay, Twist1 primarily recognized the 1500 bp upstream of the TP promoter transcription start sites (Figure 1H, left panel). To further identify the Twist1-binding sites, we reanalyzed the generic feature format data from the ChIP-seq assay using the ChIPseeker online software(Chen et al, 2014). Two individual motifs and their locations within the promoter of TP were found. Motif1 and Motif2 had the relative conserved sequences of GAGTGGCCGGTT and TGGCGTGAGC, respectively (Figure 1H, right panel). The truncated and mutated luciferase report plasmids were used for further motif identification. The TP promoter of the –1500 bp to –500 bp region was truncated to five shorter regions, in which the region3 of –1100 bp to –900 bp and region4 of – 900 bp to –700 bp contained the conserved motif (Figure 1I, upper panel). Luciferase assay results showed that all Twist1 transcriptionally activating regions contained the conserved Motif1 or Motif2 (Figure 1I, lower panel). Meanwhile, we constructed mutation plasmids with deletion mutations of Motif1 or/and Motif2. Luciferase assay results showed that Twist1 transcriptional activations were weakened when Motif1 or Motif2 was deleted. When Motif1 and Motif2 were both deleted, Twist1 transcriptional activation disappeared (Figure 1J). These results indicate that both motifs are related to the transcription of TP.

### TP stimulates pentose Warburg effect in HCC cell

RNA-seq was performed to study the potential functions of TP transcriptionally regulated by Twist1 in HCC cells. Pathway analysis results showed that Twist1 overexpression could activate the pentose phosphate pathway, glycolysis, and the metabolism of a series of carbon compounds (Figure 2A). GSEA results further proved that Twist1 promoted fructose and mannose metabolic pathways, glycolysis, the amino sugar nucleotide sugar metabolic pathway, and ATP generation from the ADP process (Figure 2B). This result suggests that the transcriptional pattern of Twist1–TP improves the energy metabolism of tumor cells. Considering that TP can promote the generation of pentose through its enzyme activity and that the deterioration of local microenvironment is an important promotion factor of tumor malignant progression by metabolic reprogramming, we speculated that TP enzyme activity affects tumor glycometabolism and further stimulates malignant progression. ELISA results showed that TP overexpression significantly promoted the secretion of TP factors in the PLC-PRF-5 cell lines and that TP knockdown significantly inhibited this secretion (Figure 2C). When Twist1 or TP was upregulated in the PLC-PRF-5 cell lines, the amount of secreted TP in the medium significantly increased. TP secretion was further increased when Twist1 and TP were both upregulated. Knockdown of TP decreased the promotion of TP secretion by Twist1. Similarly, knockdown of Twist1 or/and TP significantly inhibited the secretion of TP in Hep3B cell lines (Figure 2D).

**Figure 2.**
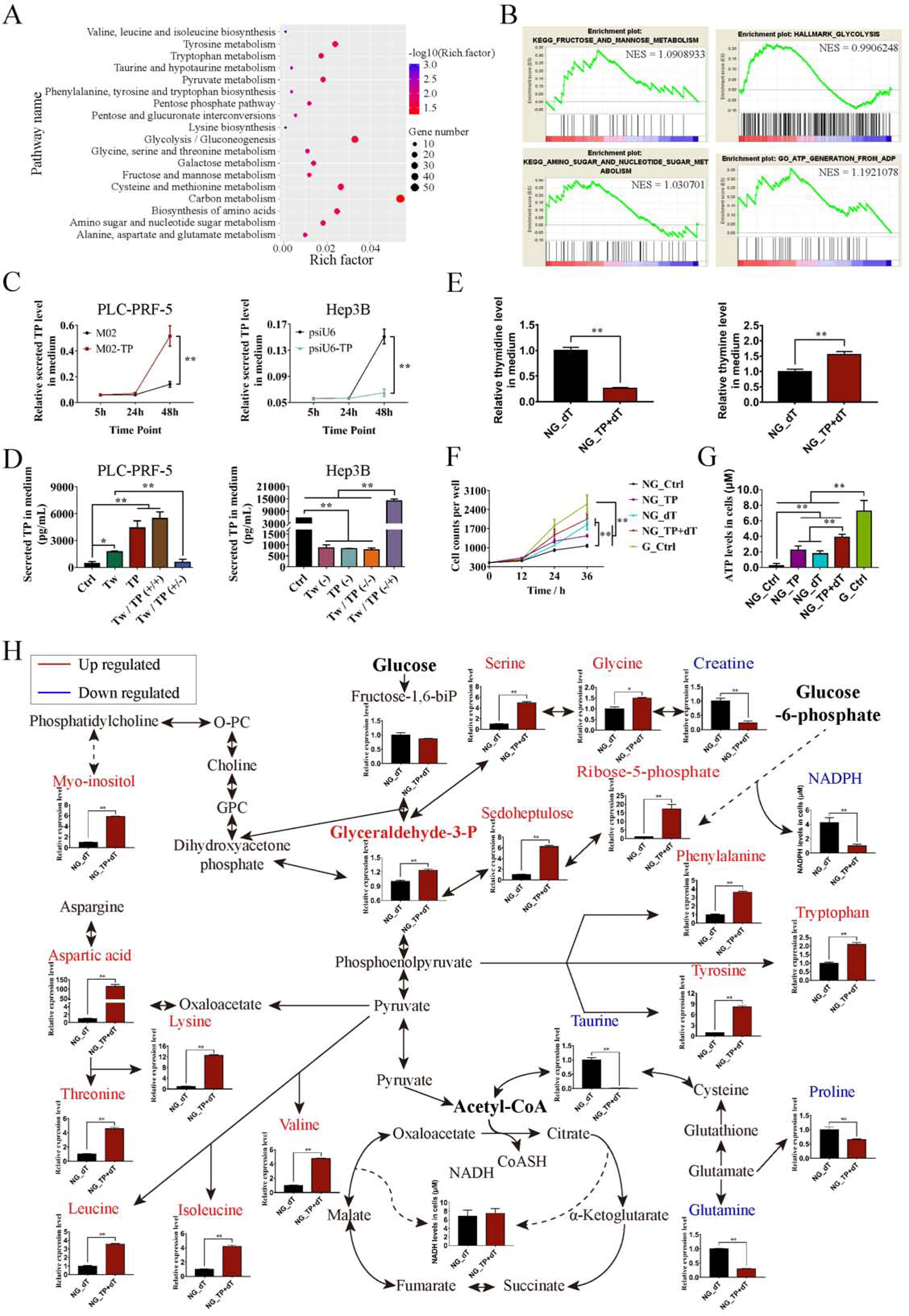
TP stimulates pentose Warburg effect in HCC cell. **A.** Overexpression of Twist1 activated a series of pathways related to glycometabolism and amino acid metabolism. **B.** Enrichment analysis of the gene expression profiles in Twist1 overexpressed PLC-PRF-5 cells by GSEA. **C.** ELISA was used to analyze the TP factor levels in medium. **D.** Correlation between Twist1/TP expressions and secreted TP factor levels in PLC-PRF-5 and Hep3B cell lines. **E.** Overexpression of TP promoted the enzymatic hydrolysis of extracellular dT in the medium. **F.** Enzymatic metabolism of extracellular dT regulated by TP promoted HCC cell proliferation. **G.** Enzymatic metabolism of extracellular dT regulated by TP promoted ATP generation in PLC-PRF-5 cell lines. **H.** Enzymatic metabolism of extracellular dT regulated by TP affected glycometabolism and amino acid metabolism in glucose-free cultured PLC-PRF-5 cell lines. (mean ± SD; n = 3 in triplicate; ^*^P < 0.05; ^**^P < 0.01)

Thymidine is a natural substrate of TP that can be catalyzed into dR-1-P and thymine. To study whether the secreted TP is an important enzyme for using thymidine as the sole carbon source of tumor cells, we added thymidine to the medium without glucose. After 24 h of incubation, the medium and PLC-PRF-5 cells were collected for further analysis. To confirm the enzyme activity of the secreted TP on extracellular thymidine, we analyzed the medium by GC–MS. Results showed that thymidine amount significantly reduced and thymine amount significantly increased in the medium (Figure 2E). Proliferation activities were detected every 12 h for a total of 36 h to study the effects of TP enzyme activity on HCC cell proliferation. The results showed that in the absence of glucose, the overexpression of TP or the addition of dT promoted PLC-PRF-5 cell proliferation, and the cell counts were further significantly increased when dT was added after TP overexpression (Figure 2F). The generation of ATP was then analyzed. Results showed that in the absence of glucose, the overexpression of TP or the addition of dT significantly promoted ATP synthesis. ATP levels were further increased when TP was upregulated with the addition of dT. When cultured with glucose-containing medium, the PLC-PRF-5 cells had the most ATP generation (Figure 2G). Further analysis results showed that in the absence of glucose, the overexpression of TP with the addition of dT did not promote the generation of NADH and NADPH. The generation of metabolic intermediates in the pentose phosphate pathway, including ribose-5-phosphate, sedoheptulose, and glyceraldehyde-3-phosphate, significantly increased. The contents of myo-inositol and amino acids derived from the glycolysis pathway, including serine, glycine, phenylalanine, tryptophan, tyrosine, aspartic acid, lysine, threonine, valine, leucine, and isoleucine, significantly increased. Meanwhile, the contents of taurine and amino acids proline and glutamine derived from TCA significantly decreased (Figure 2H). These results indicate that the main metabolic pathways affected by TP are the pentose phosphate pathway and glycolysis.

### Twist1 relied on TP to promote HCC cell migration, invasion, and VM formation through TP enzyme activities

The RNA-seq results of different treated PLC-PRF-5 cells were analyzed to further clarify the influence of pentose Warburg effect regulated by TP enzyme activities on HCC malignant progression. In the absence of glucose, the overexpression of TP or the addition of dT affected a series of pathways related to angiogenesis and metastasis, including the VEGF signaling pathway, the tight junction pathway, and the regulation of the actin cytoskeleton pathway. When TP was upregulated with the addition of dT, the influences on these pathways were more obvious (Figure 3A). The genes involved in these pathways were analyzed with PPI. Based on logFC and BetweennessCentrality index values for network nodes, the tumor VM formation-related genes KDR, FLT1, and CDH5 were revealed to be important upregulated hub genes (Figures 3B–3C). The selected PPI hubs KDR, FLT1, and CDH5 with novel connectivity exhibited potentially critical biological insights for HCC VM formation (Figure 3D). Western blot results showed that the corresponding proteins of VE-Cad, VEGFR1, and VEGFR2 were also significantly upregulated when dT was added after TP overexpression compared with the control group (Figure 3E).

**Figure 3.**
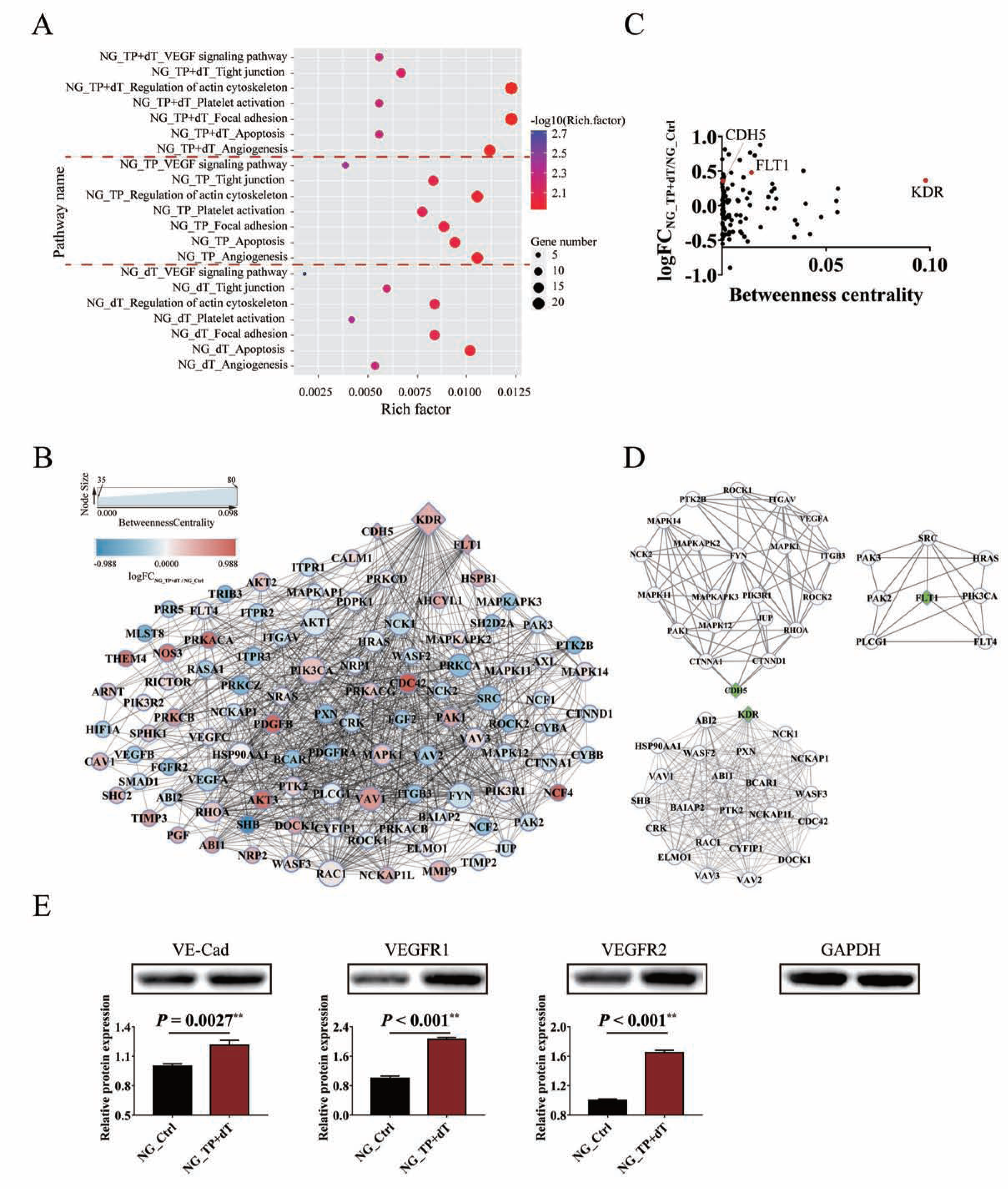
Enzymatic metabolism of extracellular dT regulated by TP affects tumor functions related to VM formation in HCC cells. **A.** Pathway analysis of glucose-free cultured PLC-PRF-5 cells affected by overexpressing TP, adding dT, or overexpressing TP and adding dT. **B.** PPI analysis of genes involved in the pathways affected by overexpressing TP and adding dT. Major hubs related to VM formation are highlighted with diamond boxes. **C.** Analysis of network topology, including logFC and BetweennessCentrality, reveals VM formation-related PPI hubs. **D.** PPI hub diagrams for the VM formation-related genes CDH5, FLT1, and KDR. **E.** Western blot was used to analyze VE-Cad, VEGFR1, and VEGFR2 expression levels influenced by overexpressing TP and adding dT in glucose-free cultured PLC-PRF-5 cell lines. (mean ± SD; n = 3 in triplicate; ^**^P < 0.01)

As VM formation was associated with cell migration and invasion, we then investigated the effect of Twist1/TP on cell migration, invasion, and VM formation in a 3D culture system in vitro by using the upregulated cell model in PLC-PRF-5 and the knockdown cell model in Hep3B. The wound healing assay showed significant differences in wound healing speed among the different transfected cells in the two cell models (Figure 4A). The invasion ability increased about four times compared with the control group when Twist1 or TP was upregulated and about six times when Twist1 and TP were both upregulated. However, when TP was knocked down, even though Twist1 was upregulated, the invasion ability of the PLC-PRF-5 cells significantly decreased. When Twist1 was knocked down and TP was overexpressed in Hep3B cells, the invasion ability was lower than that of the control group of Hep3B but significantly higher than when Twist1, TP, or both were downregulated (Figure 4B). The 3D culture assay showed that when upregulating or downregulating Twist1 or TP, the typical pipetube-like structures within the 3D Matrigel medium increased or decreased. Similarly, the network formation decreased to control level when Twist1 was upregulated and TP was knocked down in the PLC-PRF-5 cell lines. Tube formation significantly increased when the Hep3B cells were transfected with pRNAT-shTwist1 and M02-TP than when Twist1, TP, or both were knocked down in the Hep3B cells (Figure 4C). When differently treated, the PLC-PRF-5 cell lines underwent significant changes in morphology, including pseudopod disappearance and cell rounding, as detected by SEM (Figure 4D). Gradient concentrations of dT were added to the glucose-free medium of control or TP-upregulated PLC-PRF-5 cells, and the tube formations were calculated. Results showed that TP gradually promoted tube formation as dT concentration was increased from 25 μM to 100 μM (Figure 4E). To further demonstrate the relation between Twist1–TP transcriptional pattern and VM formation, we assessed the expression of VE-Cad, VEGFR1, and VEGFR2. By transfecting the luciferase reporter plasmids of VE-Cad, VEGFR1, and VEGFR2, we found that when either Twist1 or TP was upregulated in PLC-PRF-5 cells, the transcription of these three marker genes was promoted. Meanwhile, knock down of TP impaired the transcriptional activity even though Twist1 was upregulated.

**Figure 4.**
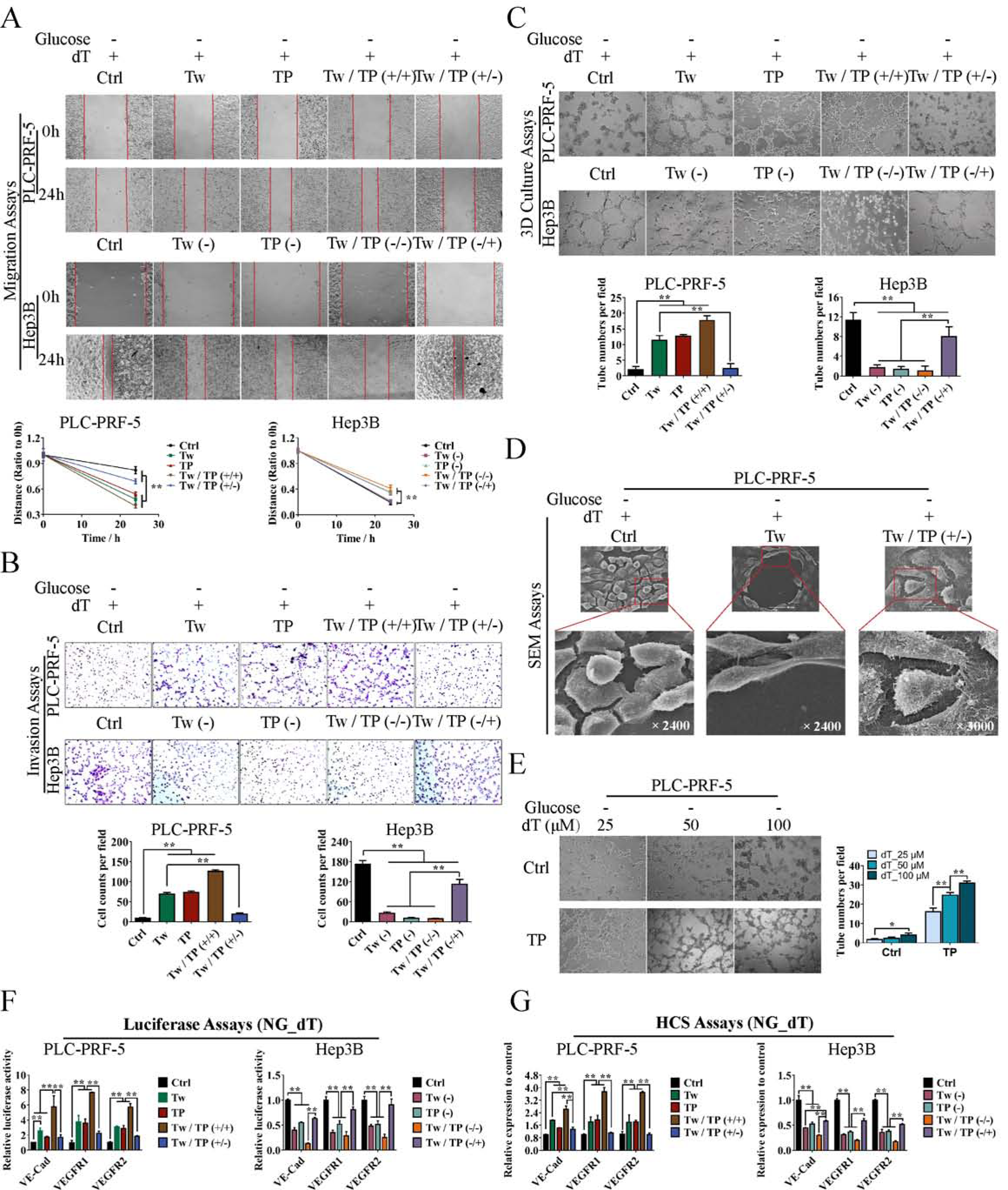
Twist1–TP transcriptional pattern promotes HCC cell migration, invasion, and VM formation through enzymatic metabolism of extracellular dT. **A.** Wound healing assay was performed. Quantitative analysis showed a significant difference in the speed of wound healing among the overexpression, knocking down, and negative control groups of glucose-free cultured PLC-PRF-5 and Hep3B cell lines. **B.** Invasion assay for different transfected PLC-PRF-5 and Hep3B cell lines cultured in glucose-free medium. **C.** In vitro assay for VM in 3D culture at 24 h. **D.** Morphological observation of the different transfected PLC-PRF-5 cell lines cultured in glucose-free media. **E.** Effects of the enzymatic metabolism of extracellular dT regulated by TP on the tube formation of glucose-free cultured PLC-PRF-5 cells. **F.** Transcriptional regulation activities of Twist1 and TP on VE-Cad, VEGFR1, and VEGFR2. **G.** Protein expression analysis of VE-Cad, VEGFR1, and VEGFR2 with the HCS systems. (mean ± SD; n = 3 in triplicate; ^*^P < 0.05; ^**^P < 0.01)

Similarly, the transcriptional activities of the marker genes significantly decreased when Twist1 and/or TP were knocked down, whereas M02-TP transfection enhanced the transcription (Figure 4F). Through the immunofluorescence assay, the three marker proteins were labeled with fluorescence and analyzed with the HCS systems. Analysis results showed that the protein expressions of VE-Cad, VEGFR1, and VEGFR2 were consistent with their transcriptional levels (Figure 4G).

### Twist1 relied on TP to promote VM formation and metastasis in vivo

To further clarify the promotion effect of the Twist1–TP transcriptional pattern on HCC VM formation and metastasis, we divided the 306 cases of clinical HCC specimens into four groups of Twist1/TP (–/–), Twist1/TP (–/+), Twist1/TP (+/–), and Twist1/TP (+/+) to analyze the correlations between Twist1/TP and HCC characteristics. Analysis results showed that when TP was negatively expressed, the malignant progression characteristics of metastasis, clinical stage, pathology stage, CEA level, AFP level, total protein level, and survival time were greatly affected even though Twist1 was positively expressed. Meanwhile, although Twist1 expression was negative in some samples, highly expressed TP significantly promoted metastasis, elevated clinical stage, promoted the protein levels of CEA and AFP, which represent the malignant progression of HCC, and reduced the survival time of HCC patients (Figures 5A–5B, Figure S2). The relation between Twist1/TP and VM formation was analyzed in our HCC clinical specimens by double labeling with CD31 and PAS. Results showed that when Twist1 and TP were both negatively expressed, VM formation was very low (0.58±0.13/HPF). VM formation increased more than 9 times when only TP was positively expressed (4.71±0.41/HPF) and nearly 12 times when both Twist1 and TP were positively expressed (8.26 ± 0.40/HPF). The specimens with Twist1 expression but without TP expression only had a VM formation of 3.07±0.24/HPF (Figure 5C). From the IHC results of the clinical HCC specimens, we found that the VE-Cad, VEGFR1, and VEGFR2 staining degrees were higher in the Twist1 or TP staining positive groups compared with those in the double negative labeling group. A much higher staining degree was observed when Twist1 and TP were both positive (Figures 5D–5E).

**Figure 5.**
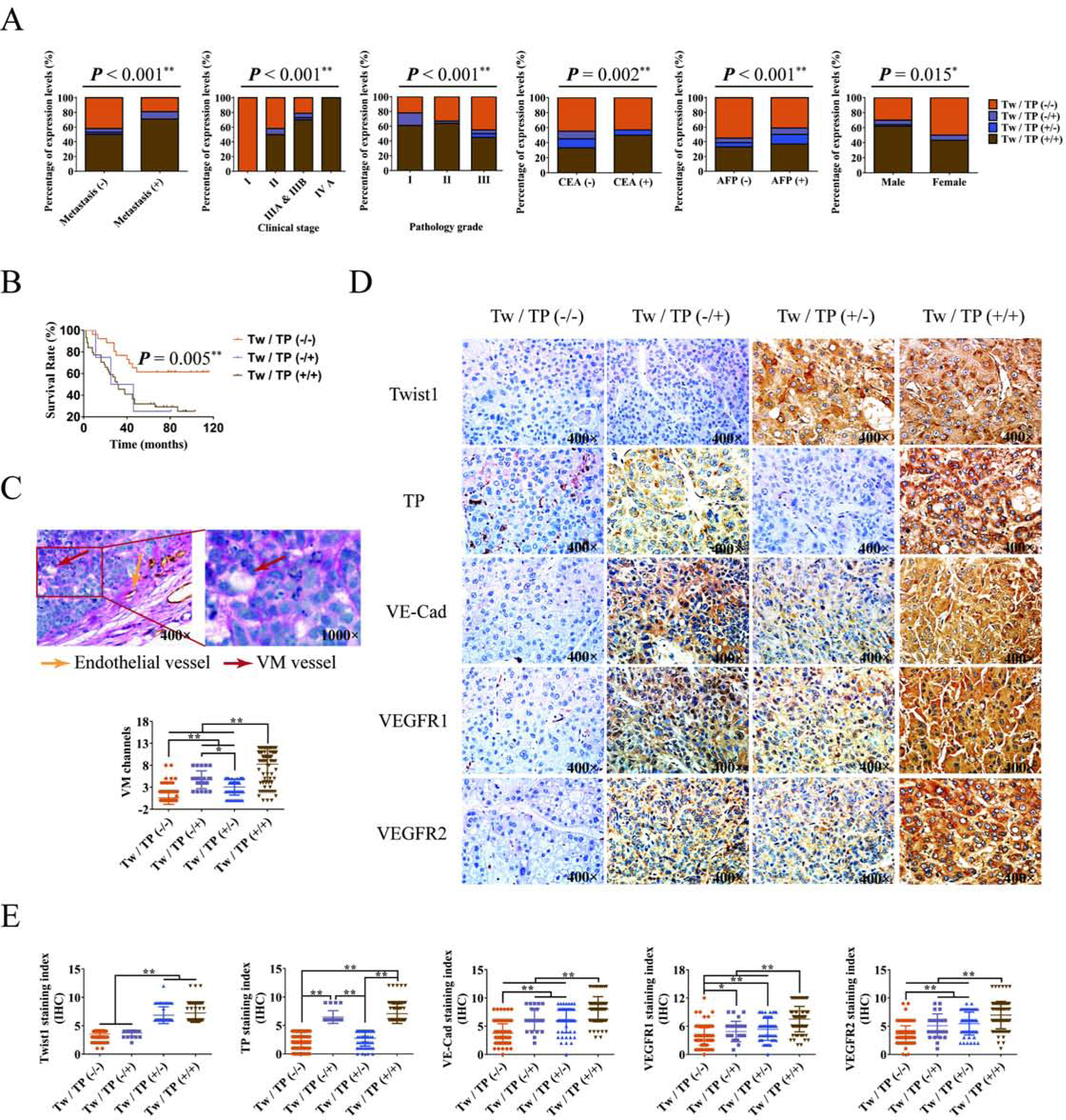
Twist1 relies on TP to promote HCC malignant progression. **A.** Correlation between Twist1/TP expressions and HCC characteristics, including metastasis, clinical stage, pathology grade, CEA level, AFP level, and gender. **B.** Overall survival analysis. **C.** Correlation analysis between VM formation and Twist1, TP expression in 306 cases of HCC. The VM channel was PAS-positive, but it did not express CD31 (red arrow). Endothelial vessel was both PAS- and CD31-positive (yellow arrow). **D.** Analysis of the HCC specimens by IHC. Samples were divided into four groups of Twist1/TP (–/–), Twist1/TP (–/+), Twist1/TP (+/–), and Twist1/TP (+/+). Results revealed that VE-Cad, VEGFR1, and VEGFR2 were minimally expressed in the Twist1 and TP negative expressed groups. When Twist1 and TP were individually or both positive, the expression levels of the three marker proteins were promoted. **E.** Statistical analysis of the protein expression in 306 HCC specimens among the four groups. (mean ± SD; ^*^P < 0.05; ^**^P < 0.01)

PLC-PRF-5 xenograft model experiments were then conducted in Balb/c-nu/nu mice. When Twist1 and TP were individually or both overexpressed, tumor growth and lung metastasis were significantly increased. However, when TP was knocked down after overexpressing Twist1, the tumor volume and lung metastasis were impaired almost to control level (Figures 6A–6B). Then, we measured VM formation through double staining with CD31 and PAS. The control group had the fewest number of VM (0.50 ±0.29/HPF) among the groups. After overexpressing Twist1 or TP, VM formations increased to 5.75±0.48/HPF and 5.50±0.65/HPF, respectively. When Twist1 and TP were both overexpressed, VM formation further increased (7.00±0.41/HPF). When Twist1 was overexpressed and TP was knocked down, VM formation was reduced to almost control level (1.50±0.29/HPF) compared with the three other groups (Figure 6C). Consistent with these results, the IHC analysis further demonstrated that Twist1 and TP promoted VM marker expression and that TP knock down weakened the promotion effect of Twist1 overexpression (Figures 6D–6E).

**Figure 6.**
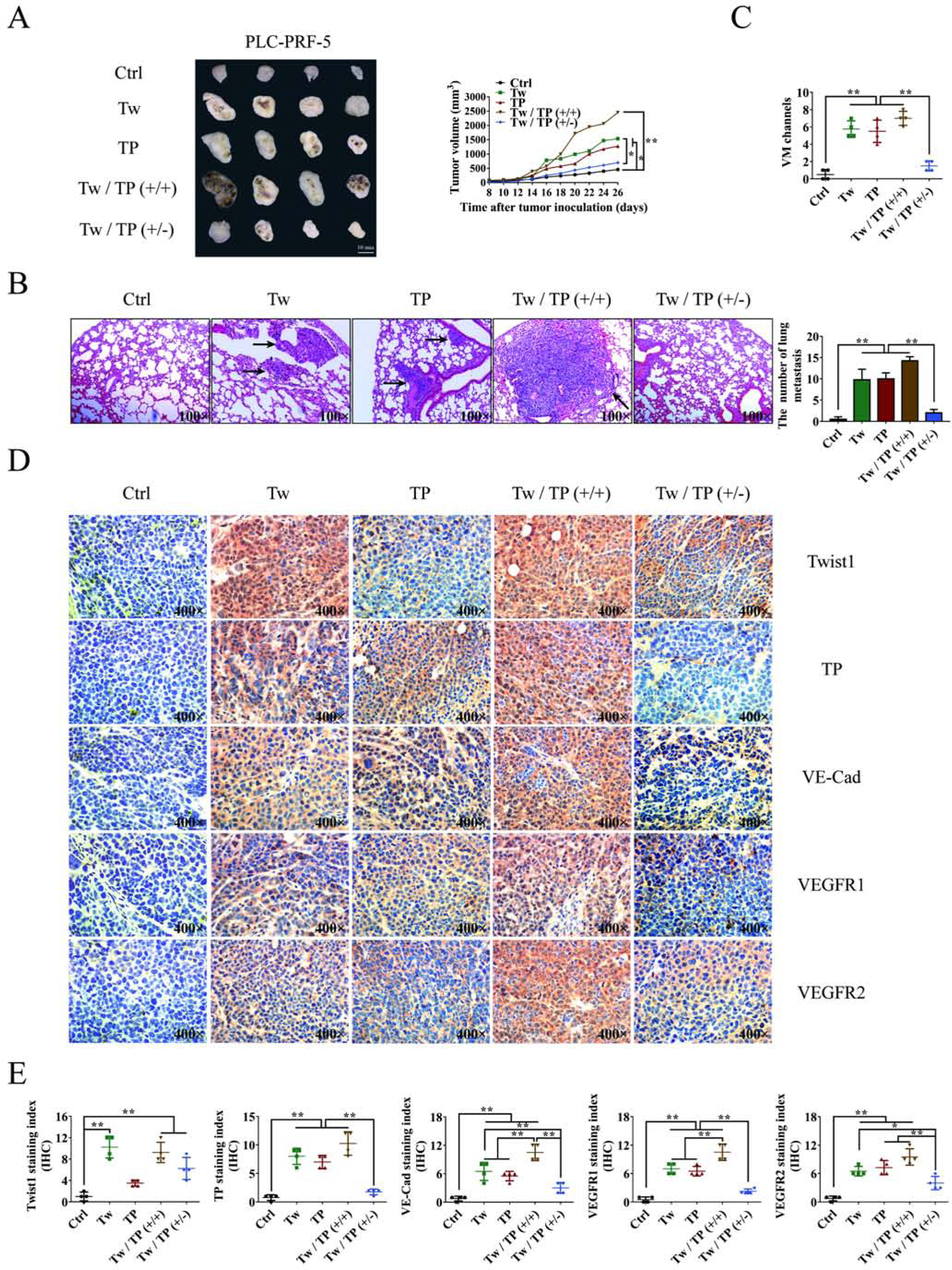
Effects of the transcriptional pattern of Twist1 to TP on HCC growth, metastasis, and VM formation in the xenograft model. **A.** Twist1 and TP promoted PLC-PRF-5 xenograft growth, and knocking down TP impaired the promotion effects of Twist1. **B.** Number of tumor formation of lung metastasis in different groups. Images were taken at 100× magnification. **C.** Statistical analysis of VM formations in different groups. **D.** Analysis of Twist1, TP, VE-Cad, VEGFR1, and VEGFR2 expression in xenograft tumors. Images were taken at 400× magnification. **E.** Statistical analysis of Twist1, TP, VE-Cad, VEGFR1, and VEGFR2 expression levels. (mean ± SD; ^*^P < 0.05; ^**^P < 0.01).

### TP enzyme inhibitor suppressed HCC VM formation and metastasis

The TP enzyme inhibitor tipiracil (TPI)(Matsushita et al, 1999) (Figure 7A) was used to clarify the promotion effects of TP on HCC VM formation and metastasis, and explore the possibility of using TP as a HCC therapy target. When TP was knocked down in the Hep3B cells, TPI had a weaker inhibitory effect on cell migration compared with the single addition of TPI (Figure 7B, Figure S3A). Similarly, knocking down TP or adding TPI significantly inhibited Hep3B cell invasion and tube formation. When TP was knocked down and then TPI was added, the suppression was not as effective as when TPI was added alone (Figures 7C–7D, Figures S3B–C). Afterward, the nude mice model transplanted with Hep3B cells was established. TPI effectively inhibited the growth of xenograft HCC. When TP was knocked down, the inhibitory effect of TPI was reduced (Figure 7E). Results showed that knocking down TP or adding TPI significantly inhibited lung metastasis, whereas knocking down TP weakened the inhibitory effect of TPI (Figure 7F, Figure S3D). Finally, the VM formation and expression of VM-related marker proteins were analyzed, and the results showed that TPI significantly inhibited VM formation and decreased the expression levels of VE-Cad, VEGFR1, and VEGFR2. When TP was knocked down, the inhibitory effect of TPI was weakened (Figures 7G–7H).

**Figure 7.**
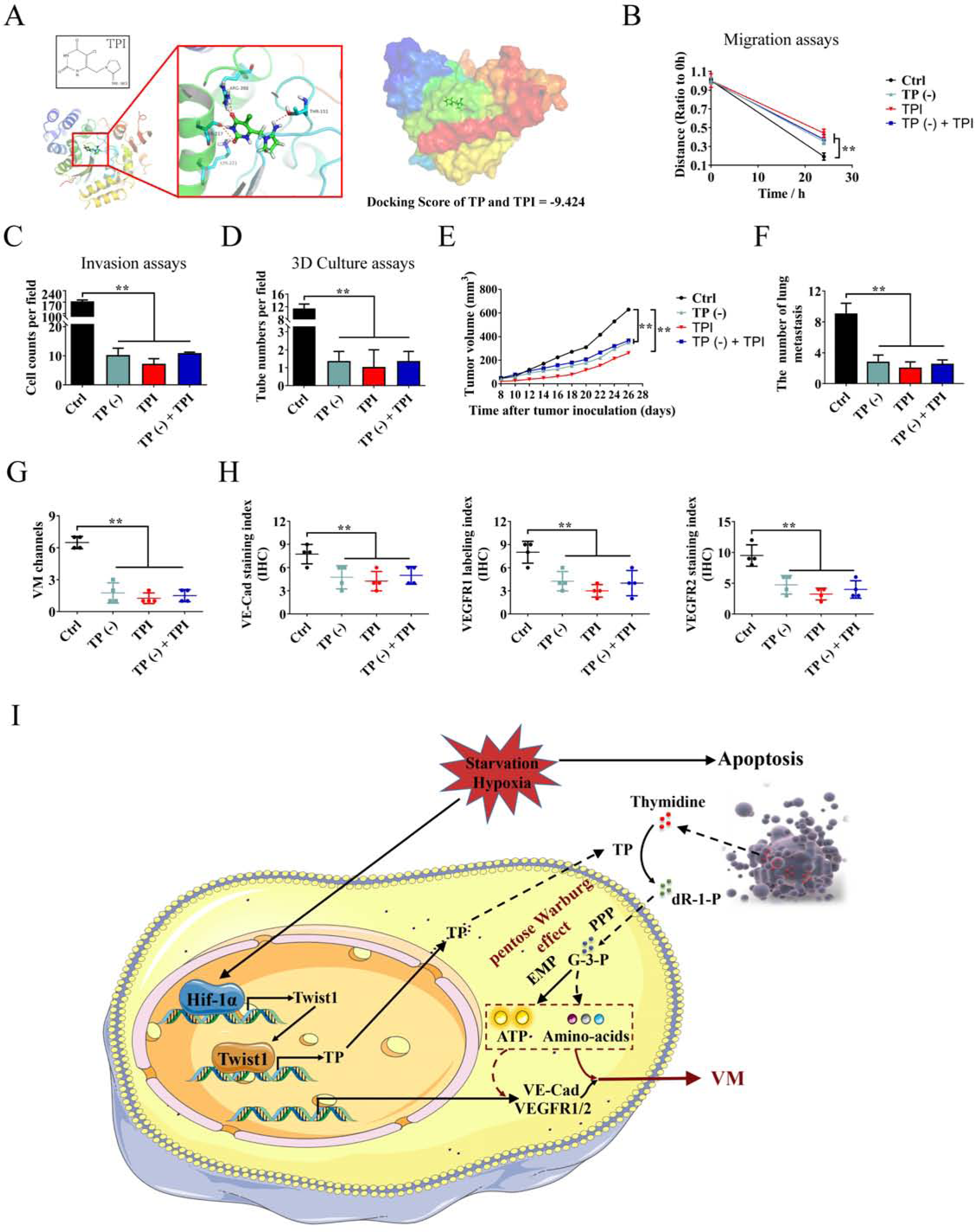
Effects of TP enzyme inhibitor on HCC VM formation and metastasis. **A.** Molecular docking results of TPI and TP. **B.** Wound healing assay was performed. Quantitative analysis showed a significant difference in the speed of wound healing among the different treated Hep3B cells. **C.** Invasion assay for Hep3B cells with knocked down TP, the addition of TPI, or knocked down TP and the addition of TPI.**D.** In vitro assay for VM in 3D culture at 24 h. **E.** Effects of knocking down TP and/or adding TPI to Hep3B xenograft tumor growth. **F.** The number of tumor formations of lung metastasis in different groups. Images were taken at 100× magnification. **G.** Statistical analysis of VM formation in different groups. **H.** Statistical analysis of VE-Cad, VEGFR1, and VEGFR2 expression levels in Hep3B xenograft tumors. **I.** The proposed regulatory mechanism between Twist1–TP and VM. (mean ± SD; ^**^P < 0.01)

## Discussion

HCC is a common malignant tumor that causes high mortality worldwide. Although hepatic resection and liver transplantation have largely improved the overall survival rate of HCC patients, distant metastasis nevertheless contributes to the failure of these treatments(Maluccio & Covey, 2012). Ischemia and hypoxia caused by the deterioration of the local tumor microenvironment are the main inducers of tumor metastasis(Doi et al, 2002; Qiu et al, 2017). VM is an effective solution to the deterioration of the local tumor microenvironment(Folberg et al, 2000). VM is accepted as a new model of neovascularization made up of aggressive tumors, and the formation of VM provides sufficient blood for tumor survival and malignant progression(Hendrix et al, 2016). However, how tumor cells deal with the deteriorated local microenvironment of hypoxia–ischemia before VM formation and whether VM formation is related to the solutions need further exploration. Hypoxia-related problems could be solved through the activated HiF-1α pathway by tumor cells(Carmeliet et al, 1998); however, the mechanism of carbon source selectivity of tumor cells is not clear during glucose deprivation caused by ischemia. Stemness and EMT are regarded as major mechanisms in VM formation and metastasis, and they are both associated with metabolic reprogramming(Jiang et al, 2015; Shen et al, 2015). Whether VM formation and metastasis are also involved in metabolic reprogramming is one of the discussion points in this study, and our results showed that metabolic reprogramming driven by TP was related to VM formation and metastasis.

Twist1, as a transcription factor, has been identified as an important EMT inducer(Martin & Cano, 2010). It binds promoter DNA sequences like the E-box motif and transcriptionally activates multiple proteins, including the VM marker of VE-Cadherin(Eckert et al, 2011; Sun et al, 2010). By TCGA data analysis and molecular assays, we demonstrated that Twist1 binds to two regions of the promoter sequences of TP and transcriptionally activates its expression. TP was first discovered in 1954 as a key enzyme of the pyrimidine salvage pathway, and in 1987, a secretory TP in peripheral blood was identified to stimulate the growth of endothelial cells(Friedkin & Roberts, 1954; Miyazono et al, 1987). Subsequently, the functions of the TP factor were associated with the enzyme activities of thymidine(Furukawa et al, 1992). Vascular supply occlusion stimulated the expression of TP, and TP was proven to induce migration and angiogenesis in both endothelial and tumor cells(Akiyama et al, 2004; Furukawa et al, 1992; Griffiths et al, 1997). By catalyzing thymidine, TP can provide pentose for tumor cells(de Bruin et al, 2003). Whether the pentose generated from TP enzyme activity is related to the deterioration of the local tumor microenvironment and whether TP is an important metabolic reprogramming protein in HCC VM formation and metastasis were investigated in this study.

By analyzing the RNA-seq and metabolomics results, we found that when glucose was free, the secreted TP can promote the generation of ATP in HCC cells and the surviving of tumor cells by catalyzing extracellular thymidine. Further analysis revealed that TP enzyme activity promoted the conversion of pentose to G-3-P. Considering the lack of glucose and no increase in NADPH content, we concluded that the upgeneration of G-3-P was not through glycolysis and the normal pentose phosphorylate pathway. Subsequently, we found that a series of amino acids and other metabolic intermediates derived from the glycolysis pathway was significantly upregulated, but the contents of NADH, taurine, and amino acids proline and glutamine derived from TCA did not increase or even significantly decreased. These results revealed that when HCC cells were faced with a deteriorated tumor local environment, TP enzyme activity caused a metabolic reprogramming of the pentose Warburg effect by coupling the pentose phosphate pathway and glycolysis. Furthermore, this metabolic reprogramming provided possibilities for HCC cell proliferation and malignant progression.

Further in vitro and in vivo experiment results showed that Twist1 relied on TP enzyme activity to promote HCC metastasis and VM formation, and the promotion effects of VM formation by TP gradually improved with increasing thymidine concentration. Mechanism study results revealed that VM formation was caused by the increase in transcription and expression of VE-Cad, VEGFR1, and VEGFR2. Considering that Twist1 can directly transcriptionally activate VE-Cad, we explored whether TP could promote the stability or expression of Twist1 or its upstream regulatory proteins such as HIF-1α(Yang et al, 2008). Unfortunately, no valid results were obtained (data not shown), which illustrated that TP influenced the expression of VE-Cad through other pathways, and it needs further explorations. Finally, TP enzyme inhibitor TPI was used, and the results further clarified the promotion effects of TP on HCC VM formation and metastasis, and demonstrated the possibility of using TP as a HCC therapy target.

In summary, this study described for the first time that TP was transcriptionally regulated by Twist1 and that TP enzyme activity promoted HCC metastasis and VM formation through pentose Warburg effect under a deteriorated tumor microenvironment (Figure 7I). This study is complementary to the theory of tumor metabolic reprogramming and further enriches knowledge on the tumor VM formation and metastasis mechanism of Twist1. Meanwhile, the study also indicated that TP might be a potential therapeutic target for HCC. Our findings might benefit further studies on the mechanism of HCC malignant progression and provide new strategies for HCC diagnosis and therapy.

## Materials and Methods

### Case selection

In this study, HCC tissue microassays containing 306 cases were purchased from US Biomax for IHC or PAS&CD31 double staining, followed by analysis of the correlation among metastasis, clinical stage, pathology grade, CEA content, AFP content, gender, survival time, VM formation, VE-Cadherin expression, VEGFR1 expression, VEGFR2 expression, Twist1 expression, and TP expression. HCC characteristics were grouped based on the best cut-off values or staining index.

### TCGA data analysis

The genomic data of cancers were downloaded from TCGA. Differentially expressed genes were screened based on a difference coefficient greater than 0.7. Afterward, co-expressed genes of Twist1 were further analyzed, and genes with a co-expression Pearson coefficient greater than 0.3 were deemed co-expressed with Twist1. The co-expressed genes of Twist1 were then Venn analyzed with the ChIP-seq results of Twist1.

### Immunohistochemical analysis

Immunohistochemistry was performed to detect the expression levels of different proteins. Tissue sections were deparaffinized in xylene and rehydrated by gradient alcohol prior to immunohistochemistry. Endogenous peroxidase activity was blocked by incubation with 3% hydrogen peroxide in methanol for 30 min. Then, the tissue sections were heated using 0.01 M citric acid buffer for 10 min in a microwave oven. Slides were stained with antibodies to Twist1 (1:100, Santa Cruz Biotechnology), TP (1:100, Abcam), VE-Cad (1:50, Abcam), VEGFR1 (1:250, Abcam), VEGFR2 (1:50, Abcam), or CD31 (1:50, Abcam). After incubation with horseradish peroxidase-conjugated goat anti-rabbit/mouse IgG, the sections were stained with 3,3’-diaminobenzidine and then counterstained with hematoxylin or periodic Acid-Schiff (PAS). Finally, the sections were dehydrated and mounted. The immunohistochemical staining index was scored by multiplying the positive degree (0 for none, 1 for weak brown, 2 for moderate brown, and 3 for strong brown) and positive rate (1 for 0–25%, 2 for 25%–50%, 3 for 50%–75%, and 4 for 75%–100%). Sections with staining indices greater than 6 were considered the high-expression group. PAS positive channels without CD31 staining were considered to be VM. Five random ×40 fields per HCC tissue were scored with a morphology consistent with mimicry. All quantification experiments were performed in a blinded setting.

### Plasmids

Plasmids of pCDNA3.1-Twist1, pRNAT-shTwist1, M02-TP, and psiU6-TP were purchased from GeneCopoeia (Guangzhou, China). The reporter gene plasmid PGL4.3-TP and its truncations and mutations were constructed by inserting the promoter regions of TP, PCR-amplified from human genomic DNA, into the PGL4.3 vector (Table S1).

### Cell culture and transfection

Human HCC cell lines SNU387, PLC-PRF-5, Hep3B, HepG2, Huh-7, MHCC-9L, and MHCC-97H were obtained from the American Type Culture Collection and KeyGen Biotech (Nanjing, China). The HCC cells were cultured in RPMI-1640 or DMEM medium (Hyclone) with 10% fetal bovine serum (Hyclone). Glucose-free media RPMI-1640 (Life Technologies, China) and DMEM (Neuronbc, China) were used for the starvation experiments, and 100 μM thymidine (Amresco, USA) was added for further incubation. TPI (Meilunbio, China) at 10 μM was added to the medium for enzyme inhibition experiments. The vectors were transfected into cells using the Roche transfection reagent (Roche, Switzerland).

### Western blot analysis

Immunoblots were performed with samples that contain total protein (30 μg) through SDS-PAGE and then transferred onto PVDF membranes (Millipore, USA). The membrane was incubated with the primary antibody of Twist1 (Santa Cruz Biotechnology, USA), thymidine phosphorylase (Abcam, UN), or GAPDH (Affinity Bioreagents, USA), followed by incubation with the second antibody (Santa Cruz Biotechnology, USA). Blots were detected by using an enhanced chemiluminescence detection kit (Millipore, USA). Densitometric analysis was performed using the ImageJ software.

### Luciferase assays

After transfection of the reporter gene plasmids, the cells were cultured for another 48 h. Transactivation assays were performed with Dual-Luciferase Assays System (Promega), and luciferase activities were measured using the Luminoskan Ascent reader system (Thermo Scientific, USA).

### RNA-seq and omics analysis

PLC-PRF-5 cells transfected with control plasmids, pCDNA3.1-Twist1, or M02-TP; added with thymidine (dT); or added with dT after transfection of M02-TP were collected with TRIzol (Sigma, USA) for further RNA isolation. After preparation of the biotinylated complementary RNA (cRNA) using the Illumina TotalPrep RNA Amplification Kit (Ambion), the cRNA was hybridized with BeadChip. Samples were prepared as technical duplicates. Data were acquired with Illumina BeadChip Reader and evaluated using the Illumina BeadStudio Application by Genergy Bio (China). Gene set enrichment analysis (GSEA)(Subramanian et al, 2005) and Gene ontology (GO) analysis were used to detect changes in biological process, cellular components, and molecular function. Pathway analysis was conducted based on the KEGG database. Protein–protein interaction (PPI) was analyzed using the STRING database(Szklarczyk et al, 2015) and Cytoscape software (NRNB)(Bader & Hogue, 2003).

### ELISA

The commercial ELISA kit of TP (R&D, USA) was used to measure the TP concentrations in the culture medium in accordance with the manufacturer’s instructions. Each experiment was repeated in triplicate, and the mean values (mean ± SD) were presented.

### Metabolite detection

The metabolite of ATP was detected using the ATP Assay Kit (Beyotime, China). Metabolites of NADH and NADPH were detected using the NAD^+^/NADH and NADP^+^/NADPH assay kits (BioAssay Systems, USA). Other metabolites in the PLC-PRF-5 cells treated with dT and overexpressed with or without TP were detected by GC-TOF/MS.

### Wound healing assay

The transfected cells were seeded into 24-well culture plates at a density of 5 × 10^5^ cells/well. Tumor cells adhering to the plate were removed by scratching the wound with a 200 µL pipette tip and then incubated with serum-free medium for 24 h. Images of the wounds were taken under a light microscope (Nikon, Japan). Each experiment was repeated in triplicate, and the mean ± SD were presented.

### Matrigel invasion assay

Cell invasion assay was performed using Transwell cell culture inserts (Corning, USA) and Matrigel (BD, USA). The transfected cells were allowed to invade for 24 h. The passed cells were fixed in 4% paraformaldehyde, stained with crystal violet solution, and then counted under a light microscope (Nikon, Japan).

### Three-dimensional culture assay

PLC-PRF-5 and Hep3B cells transfected with different plasmids were seeded into a 48-well plate pre-coated with Matrigel and then cultured with 250 µL RPMI-1640 or DMEM medium containing 10% FBS for 24 h. Images of the tube were taken under a light microscope (Nikon, Japan). Other samples were seeded on climbing slides; fixed with glutaraldehyde, osmic acid, and tert-butyl alcohol; coated with a thin layer of gold; and then imaged under a scanning electron microscope (SEM, LEO 1503 VP, Germany).

### Immunofluorescence staining

The transfected cells were plated into 96-well plates at a density of 4000 cells/well. After fixed with ice-cold methanol, the cells were blocked with serum. The primary antibodies VEGFR1, VEGFR2, and VE-Cad (Abcam, USA) were used at a 1:100 work solution. Fluorescein isothiocyanate- and tetramethyl rhodamine isothiocyanate-conjugated mouse and rabbit immunoglobulin G antibodies (Santa Cruz Biotechnology, USA) were used to label immunofluorescence. After immunolabeling, the cells were stained with Hoechst (Sigma, Germany) and then measured with high-content screening (HCS) systems (Thermo Scientific, USA).

### Molecular docking

Molecular docking was performed using Schrodinger software. The crystal structure of TP was downloaded from the PDB database (https://www.rcsb.org, PDB code: 2WK6). The ligand in TP was extracted from the crystal structure as the location center of docking. The structure of TPI was generated by Chemdraw and optimized by the LigPrep module of Schrodinger. The Glide XP (extra precision) mode was used for docking score calculation.

### Animal studies

Twenty BALB/c nu/nu mice that were 5 to 6 weeks old, with an equal amount of males and females, were divided randomly into five groups. PLC-PRF-5 cells transfected with control plasmids, pCDNA3.1-Twist1, M02-TP, pCDNA3.1-Twist1 and M02-TP, or pCDNA3.1-Twist1 and psiU6-TP were subcutaneously injected into the flank in separate groups at a density of 5 × 10^6^ cells/mouse. After tumor inoculation, the tumor size was measured every 2 days for another 18 days. In the enzyme inhibition experiments, the tumor-bearing mice were treated with TPI at a dose of 45 mg/kg/day orally every 2 days for 26 days. Subsequently, all mice were sacrificed, and xenografts and lungs were collected. Lung tissues were harvested for histologic examination to measure the metastasis, and the xenografts were further analyzed by IHC and PAS&CD31 double staining.

### Statistical analysis

Differences among the groups were analyzed by one-way ANOVA using the Bonferroni post hoc test. Qualitative variables were analyzed by Fisher’s exact two-tailed test or the chi-squared test. Overall survival analysis was performed using the Kaplan–Meier method with the log-rank test. All data in the study were evaluated with IBM SPSS Statistics 22.0 software (Chicago, IL, USA). *P* values less than 0.05 were deemed statistically significant.

## Acknowledgments

This study was supported by the National Natural Science Funds of China [Grants 81572838, 81402973 and 81703581]; Tianjin Natural Science and Technology Fund [Grant 15JCYBJC26400]; National Science and Technology Major Project [Grant 2017ZX09306007]; and Tianjin Science and Technology Project [Grant 15PTGCCX00140].

## Author Contributions

C.Y., T.S., Y.-y.L., Q.Z., Y.Q., and J.-m.Z. are responsible for this study. T.S. conceived and designed the study. T.S. and Q.Z. proposed the concept of abnormal pentose metabolic pathway. C.Y. revised the paper and provided financial support. Y.-h.T. carried out the bioinformatics analysis of TCGA and RNA-seq with assistance from W.-l.Z. X.-j.H. performed the metabolic detection experiment. Y.Q., M.-m.L., and S.-m.Z. assisted with the animal experiments. J.-m.Z., M.L., H.-l.T., Z.Z., S.C., H.-j.L., L.Y. and H.-g.Z. performed all other experiments. Q.Z., Y.Q. and J.-m.Z. analyzed data. Q.Z., Y.-y.L., T.S., and C.Y. wrote the manuscript.

## The Paper Explained

### Problem

Metastasis and vasculogenic mimicry (VM) are typical tumor progression, which is dependent on metabolic reprogramming. Thymidine phosphorylase (TP), as a phosphorylase, catalyzes thymidine to generate pentose. In our study, we found that Twist1, a tumor progression inducer, transcriptionally regulated TP. It remains unclear whether TP could promote tumor progression by metabolic reprogramming.

### Results

Here, we show that TP activates extracellular thymidine metabolization into ATP and amino acids through pentose Warburg effect by coupling the pentose phosphate pathway and glycolysis. And Twist1 relies on TP-induced metabolic reprogramming to promote HCC VM formation.

### Impact

The findings clarify that TP plays a critical role in the HCC progression through activated pentose Warburg effect. Targeting TP may provide new strategies for HCC diagnosis and therapy.

